# E3 ligase recruitment by UBQLN2 protects substrates from proteasomal degradation

**DOI:** 10.1101/2024.07.04.602059

**Authors:** Ashley N. Scheutzow, Sachini Thanthirige, Gracie Siffer, Alexander D. Sorkin, Matthew L. Wohlever

**Author notes:** These authors contributed equally (Ashley N. Scheutzow, Sachini Thanthirige).

## Abstract

Ubiquilins are a family of proteins critical to cellular proteostasis that are also linked to several neurodegenerative diseases, with specific mutations in UBQLN2 causing dominant, X-linked ALS. Despite an initial characterization as proteasomal shuttle factors, Ubiquilins have paradoxically been reported to stabilize numerous substrates. The basis of this triage decision remains enigmatic. Many other fundamental aspects of Ubiquilin function are unclear at the mechanistic level, such as the physiological significance of Ubiquilin phase separation, the unique role of each Ubiquilin paralog, and the mechanistic defects of ALS mutants. To address these questions, we utilized a library of triple knockout (TKO) rescue cell lines with physiological expression of single Ubiquilin paralogs or disease mutants in an isogenic background. Our findings reveal that UBQLN2 has a unique ability to protect substrates from degradation and that substrate stabilization correlates with the recruitment of multiple E3 ligases, including SCF^bxo7^. We propose that E3 ligase recruitment promotes UBQLN2 phase separation, which protects substrates from proteasomal degradation. Consistent with this model, we demonstrate that ALS mutants, which were previously shown to have altered phase separation properties, also show a defect in substrate stabilization. Finally, we show that substrate stabilization appears to be a general feature of proteins that interact with the UBQLN2 Sti1 domains as amyloid precursor protein (APP) is also protected from proteasomal degradation by the formation of biomolecular condensates. This proposal unifies many existing observations in the field and presents a new paradigm for understanding Ubiquilin function in neurodegenerative disease.

## Introduction

Protein homeostasis (proteostasis) involves a complex network of cellular pathways responsible for the synthesis, folding, and degradation of proteins^1–5^. Failures in this essential process lead to numerous neurodegenerative diseases, including Amyotrophic Lateral Sclerosis (ALS)^6–12^. Ubiquilins are a family of proteins at the nexus of the cellular proteostasis network, with connections to the ubiquitin proteasome system, autophagy, and the pathological protein aggregates characteristic of Alzheimer’s disease and numerous other proteinopathies^13–18^. Humans have three widely expressed Ubiquilin paralogs, UBQLN1, UBQLN2, and UBQLN4. The unique role of each paralog is poorly understood, although UBQLN2 has garnered significant attention as point mutations lead to dominant, X-linked ALS^19–25^. These mutations lead to the formation of protein aggregates that are a hallmark of ALS pathology, but the precise mechanisms by which UBQLN2 mutations contribute to ALS remain enigmatic^26–29^.

Ubiquilins were originally characterized as proteasomal shuttle factors that facilitate protein degradation, with the Ubiquitin Like (UBL) domain binding the proteasome and the Ubiquitin Associated (UBA) domain binding to ubiquitinated proteins^30,31^. The middle region of the protein contains two Sti1 domains which are important for binding to hydrophobic transmembrane domains (TMDs)^32^. Several groups have previously shown that the UBA domain can also recruit E3 ligases, although the functional role of E3 ligase recruitment is unknown^32,33^. Despite the initial characterization as shuttle factors and the association with E3 ligases, Ubiquilins have paradoxically been shown to stabilize numerous substrates^29,34–39^. Why some substrates are stabilized and others are degraded is unknown.

Another defining feature of Ubiquilins is the ability to undergo phase separation and form biomolecular condensates^13^. The role of phase separation in Ubiquilin activity is unclear, although there are hints that it is linked to the substrate triage decision. For example, Ubiquilin phase separation is modulated by polyubiquitin chains in a linkage dependent manner and proteasome activity can be inhibited in Ubiquilin condensates^40,41^. Interestingly, many ALS mutants show altered phase separation properties^23,26,42^.

Here, we use a library of triple knockout (TKO) rescue cell lines that have physiological expression of a single Ubiquilin paralog or disease mutant in an isogenic background^32^. Using this library, we show that TKO cells rescued with WT UBQLN2 have a unique ability to protect substrates from proteasomal degradation whereas rescue with ALS causing mutations leads to reduced levels of substrate stabilization. Interestingly, substrate stabilization correlates with UBQLN2 expression levels, E3 ligase recruitment, and the formation of biomolecular condensates. Building off recent work demonstrating that polyubiquitin chains promote UBQLN2 phase separation^43–45^, we propose that E3 ligase recruitment leads to the formation of biomolecular condensates which protect substrates from proteasomal degradation.

## Results

### Cells expressing only UBQLN2 show growth defects

To examine the unique functional role of the different Ubiquilin paralogs, we used the TRex Flp-In system to generate triple knockout rescue HEK cell lines **(Figure S1A)**. The triple knockout (TKO) cells harbor deletions UBQLN1, 2, and 4 and are then complemented with a single Myc-tagged Ubiquilin variant that is chromosomally integrated and under control of a doxycycline inducible promoter^32^. We will refer to the rescue cell lines as TKO^UBQLN_^, with the superscript describing the Ubiquilin variant used for complementation. Importantly, all integrations occur at the same chromosomal locus, so all cell lines are isogenic except for the Ubiquilin of interest.

As Ubiquilin phase separation is concentration dependent, care must be taken to replicate physiological Ubiquilin expression levels. We first sought to define the concentration of doxycycline required for physiological expression. Using anti-UBQLN2 and anti-UBQLN4 antibodies, we determined that 5 ng/mL doxycycline gave physiological expression levels of UBQLN1 and UBQLN4 whereas 0.4 ng/mL doxycycline was required to give physiological expression of UBQLN2 **(Figures S1B-D)**. As many previous studies used transient overexpression of UBQLN2, we decided to study rescue cells expressing UBQLN2 at physiological levels (0.4 ng/mL) and overexpression conditions (5 ng/mL), which we will refer to as TKO^UBQLN2^ and TKO^UBQLN2-High^, respectively.

With our expression conditions firmly established, we first examined the fitness of the TKO rescue cells. Consistent with previous reports^32^, TKO cells showed a significant growth defect whereas TKO^UBQLN1^ and TKO^UBQLN4^ cells showed little to no fitness defect under standard growth conditions. Surprisingly, TKO^UBQLN2^ cells showed an intermediate phenotype between WT HEK cells and TKO cells **(Figure S1E)**. This effect was not dependent on UBQLN2 overexpression as TKO^UBQLN2-High^ exhibited the same growth rate.

### UBQLN2 protects client proteins from proteasomal degradation

We next asked if the growth defect in the TKO^UBQLN2^ cell line was due to changes in proteostasis capacity. As a model substrate, we used mitochondrial membrane protein ATP5G1 with the mitochondrial targeting sequence deleted (ATP5G1ΔMTS), as this protein was previously shown to bind to Ubiquilin Sti1 domains^32^. To assess the ability of Ubiquilins to facilitate degradation of ATP5G1ΔMTS we used a ratiometric fluorescent reporter containing an in frame-fusion of GFP-ATP5G1ΔMTS and mCherry separated by the viral 2A sequence^46^ **(Figure 1A)**. GFP-ATP5G1ΔMTS and mCherry are expressed at equal levels, but as separate protein products, thus providing an internal expression control. The GFP:mCherry ratio in each cell can be quantitatively measured by flow cytometry. Control experiments with untagged GFP showed nearly identical GFP/mCherry ratios between all cell lines, regardless of Ubiquilin expression **(Figure S2)**.

**Figure 1:**
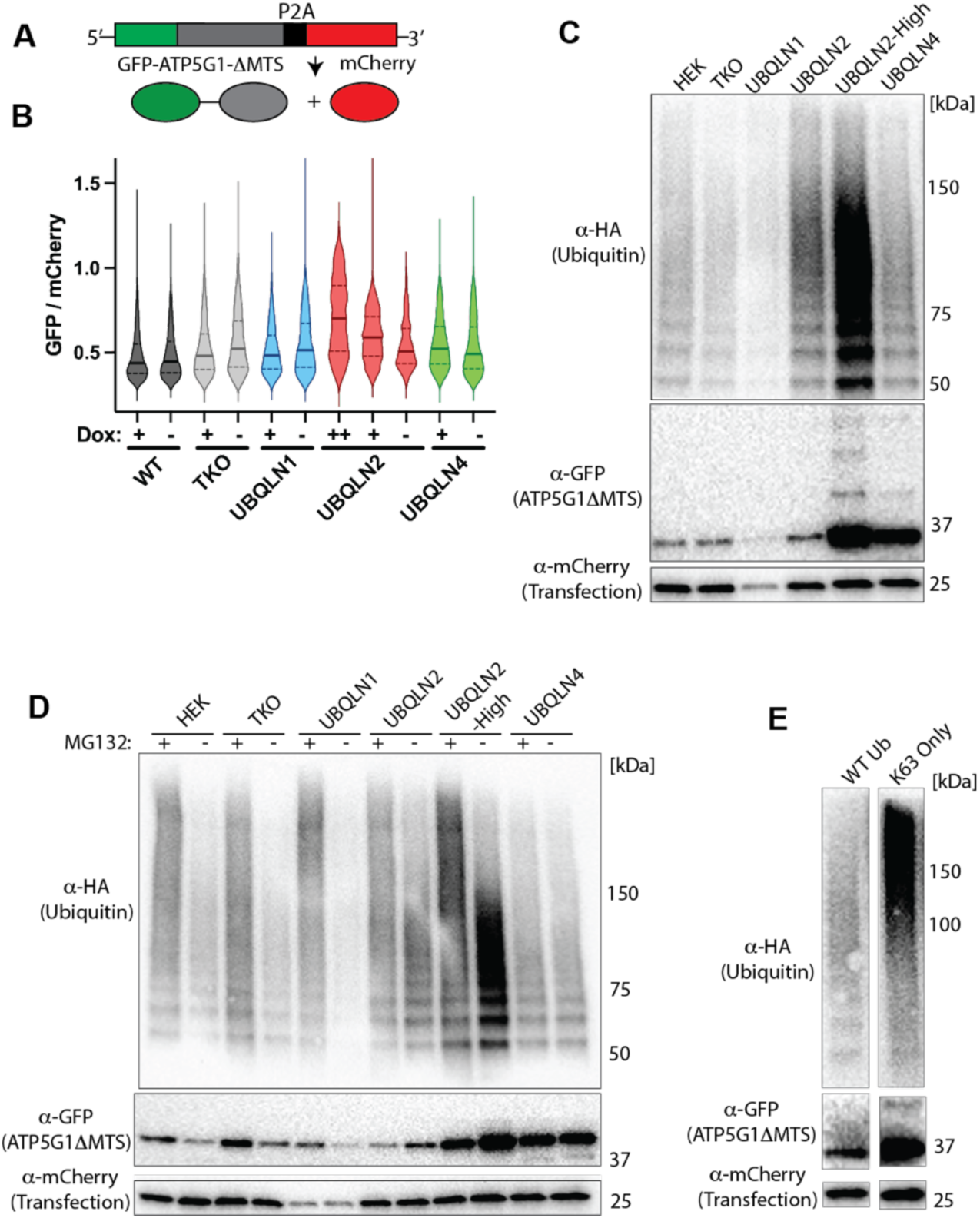
UBQLN2 protects substrates from proteasomal degradation. A) Diagram of the GFP-mCherry assay B) TKO^UBQLN2^ cells (red) show a dramatic stabilization of the substrate that correlates with UBQLN2 expression levels. Doxycycline concentrations were sufficient to give physiological Ubiquilin expression (+) or overexpression (++). Note that the GFP/mCherry ratio is much higher than in the TKO cells, suggesting that this is not a simple loss of function. C) TKO^UBQLN2^ cells have an accumulation of ubiquitinated substrates that correlates with UBQLN2 expression levels. The GFP-ATP5G1ΔMTS-P2A-mCherry substrate was co-transfected with HA-Ubiquitin. GFP immunoprecipitation shows an accumulation of Ubiquitinated substrates in TKO^UBQLN2^ cells. D) Proteasome inhibition does not increase the amount of ubiquitinated substrate in TKO^UBQLN2^ cells. Assay performed as in C, except cells were treated with MG132 or DMSO for 4 hours prior to harvesting. E) K63-only ubiquitin promotes the accumulation of ubiquitinated substrates. Cell-based ubiquitination assay with TKO^UBQLN2^ cells performed as in C except WT or K63-only HA-ubiquitin. Images were cropped from the same western with the same contrast adjustment.

We transfected the GFP-ATP5G1ΔMTS-P2A-mCherry construct in WT, TKO, and TKO rescue cell lines. The GFP:mCherry ratio decreased from ∼1 to ∼0.5 in the TKO cells, suggesting Ubiquilin independent degradation **(Figure 1B)**. We observed modest, but reproducible decreases in the GFP/mCherry ratio in WT cells compared to TKO cells, suggesting that Ubiquilins facilitate ATP5G1ΔMTS degradation, consistent with previous studies^32^. The GFP/mCherry ratio in TKO^UBQLN1^ was similar to TKO cells whereas the ratio was slightly elevated in TKO^UBQLN4^ cells.

Strikingly, we observed that ATP5G1ΔMTS was stabilized in the TKO^UBQLN2^ cell line with the degree of substrate stabilization correlating with UBQLN2 expression levels **(Figure 1B)**. Because the overall increase in GFP:mCherry ratio far exceeds the level in TKO cells, we conclude that UBQLN2 is actively protecting substrates from degradation rather than acting via a simple loss of function.

This prompted us to investigate how UBQLN2 can protect substrates from degradation. As Ubiquilins can bind to the proteasome, we hypothesized that UBQLN2 could be protecting client proteins from proteasomal degradation. Indeed, Valentino et al. recently reported that UBQLN2 can recruit the 26S proteasome into biomolecular condensates, which can inhibit proteasome activity of K63 linked or mixed-linkage substrates^40^. Such a model makes several predictions, including: 1) there should be a unique accumulation of ubiquitinated substrates in TKO^UBQLN2^cells, 2) proteasome inhibitors should produce no further substrate stabilization, and 3) K63 ubiquitin linkages will enhance substrate stabilization.

To query the ubiquitination state of the substrates, we performed a cell-based ubiquitination assay. TKO rescue cell lines were co-transfected with the GFP-ATP5G1ΔMTS-P2A-mCherry and HA-Ubiquitin. GFP-ATP5G1ΔMTS was immunoprecipitated in the absence of any proteasome inhibitors. We observed a hyperaccumulation of ubiquitinated substrate only in TKO^UBQLN2^ cells, with the degree of accumulation correlating with the level of UBQLN2 expression **(Figure 1C)**.

We then repeated the cell-based ubiquitination assay in cells treated with the proteasome inhibitor MG132 **(Figure 1D)**. Proteasome inhibition increased the accumulation of both unmodified and ubiquitinated substrates in HEK, TKO, and TKO^UBQLN1^ cell lines, suggesting that ATP5G1ΔMTS is subject to proteasomal degradation in these cell lines. Conversely, addition of MG132 to TKO^UBQLN2^ cells led to a decrease in the amount of unmodified substrate. The overall amount of ubiquitinated substrates appears relatively constant, but there is a notable shift to longer polyubiquitin chains on the substrate.

To test if K63 ubiquitin linkages enhance substrate stabilization, we repeated the cell-based ubiquitination assay with TKO^UBQLN2^ cells transfected with WT or K63-only HA-Ubiquitin. Consistent with our model, we saw an increase in ubiquitinated substrates when cells were transfected with K63-only ubiquitin compared to wild type ubiquitin **(Figure 1E)**. We conclude that UBQLN2 can actively protect ubiquitinated substrates from proteasomal degradation.

### UBQLN2 has a unique ability to recruit multiple E3 ligases

We next asked what is unique about UBQLN2 that allows it to protect substrates from proteasomal degradation. Previous reports had demonstrated that UBQLN2 condensates can inhibit the 26S proteasome and UBQLN2 phase separation is enhanced by polyubiquitin chains^40,45,47,48^. Furthermore, we and others had previously demonstrated that Ubiquilins use the UBA domain to recruit an E3 ligase^32,33^. We therefore hypothesized that the accumulation of ubiquitinated substrates in TKO^UBQLN2^ cells was due to enhanced E3 ligase recruitment by UBQLN2.

To test this hypothesis, we performed a previously established *in vitro* ubiquitination assay^32^ **(Figure 2A)**. In this assay, the known Ubiquilin substrate Omp25 is generated by in vitro translation (IVT) in rabbit reticulocyte lysate (RRL) supplemented with physiological concentrations of flag-tagged Ubiquilin. Previous results had demonstrated that the UBL domain is a negative regulator of E3 ligase recruitment, so the ΔUBL construct was used as a positive control and ΔUBA as a negative control^32^. Ubiquilin substrate complexes were purified by anti-flag immunoprecipitation and the eluate was complemented with purified E1 (Ube1), E2 (UbcH5A), Ubiquitin, and ATP. Ubiquitination activity was analyzed by anti-ubiquitin western blot. As no E3 ligase was added, any ubiquitination activity must result from an E3 ligase that co-immunoprecipitated with Ubiquilin. Consistent with our hypothesis, UBQLN2 constructs had the highest level of ubiquitination activity **(Figure 2B & S3)**. We conclude that UBQLN2 has an enhanced ability to recruit E3 ligases, which correlates with substrate stabilization.

**Figure 2:**
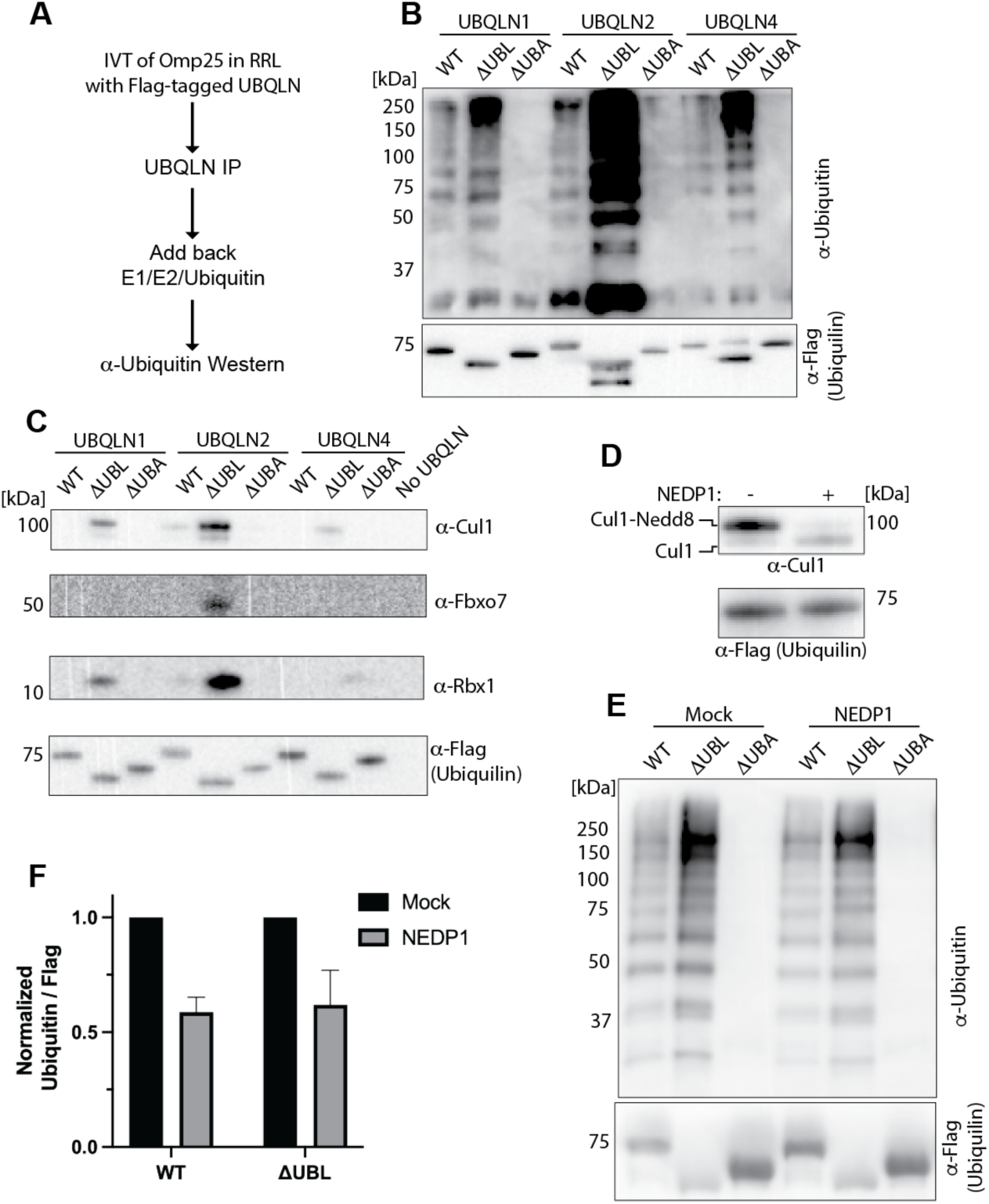
Substrate stabilization correlates with E3 ligase recruitment. A) Workflow for the *in vitro* ubiquitination assay. B) *In vitro* ubiquitination assay shows that UBQLN2 has more robust E3 ligase recruitment than UBQLN1 and UBQLN4. The ΔUBL constructs were used as a positive control and ΔUBA constructs as a negative control. Blot is overexposed to demonstrate that full-length Ubiquilin can recruit E3 ligases. See supplemental Figure S3 for additional replicates of the assay. C) UBQLN2 has a unique ability to robustly interact with components of the E3 ligase SCF^bxo7^. All blots are from the same IP. D) NEDP1 treatment of RRL prior to anti-Flag IP leads to deneddylation of Cul1 and an overall decrease in the amount of Cul1 recruited by Ubiquilin. IP was performed with ΔUBL UBQLN1. E) Treatment of RRL with NEDP1 leads to a drop in ubiquitination activity. Assay performed as in C, except RRL was treated with NEDP1 before anti-Flag IP. F) Quantification of blots in E (n = 3). Y axis is intensity of anti-ubiquitin blot divided by intensity of anti-flag blot. Intensity is normalized to mock treated lanes.

### Ubiquilins recruit activated SCF^bxo7^

We next asked which E3 ligase(s) are recruited by Ubiquilins. As previous work had shown that UBQLN1 recruits E3 ligases^32,33^, we added recombinantly purified, flag-tagged full-length or ΔUBL UBQLN1 to RRL and performed an anti-flag IP. We then submitted the elution for mass spectrometry analysis **(Tables S1 & S2)**. Our data set contained several E3 ligases, including Ube3A/E6AP, which was recently shown to interact with the Ubiquilin UBA domain^33^. Consistent with the weak 460 μM K_D_ of this interaction^33^, we were unable to detect interaction via immunoprecipitation using the roughly physiological concentration of 10 μM Ubiquilin as an input. However, upon increasing the input concentration of ΔUBL UBQLN2 to 30 μM, we were able to validate the interaction with Herc1 **(Figures S3C)**.

We also observed Cul1 and the F-box protein Fbxo7 in our mass spec results, suggesting that Ubiquilins may also recruit SCF^bxo7^. We again performed an IP with physiological levels (10 μM) of flag-tagged Ubiquilin added to RRL. We were able to validate Ubiquilin dependent recruitment of SCF components Cul1, Fbxo7 and Rbx1. Consistent with the results of the ubiquitination assay, UBQLN2 had the most robust recruitment of SCF^bxo7^ components **(Figure 2C)**.

We hypothesized that if SCF^bxo7^ is the major E3 ligase recruited by Ubiquilins, then removal of Nedd8 from Cul1 should dramatically reduce ubiquitination activity since SCF^bxo7^ activity requires neddylation of Cul1^49^. To test this hypothesis, we added the Nedd8 specific protease NEDP1 to the RRL prior to performing the anti-Flag IP with either full-length or ΔUBL UBQLN1. As expected, addition of NEDP1 collapsed almost the entire Cul1 signal down to a lower molecular weight band, consistent with de-Neddylation activity **(Figure 2D)**. Interestingly, treatment with NEDP1 also leads to a substantial decrease in the amount of Cul1 bound to Ubiquilin, suggesting that Ubiquilins preferentially recruit the activated form of the enzyme. We then performed our ubiquitination assay with NEDP1 treated RRL. Treatment with NEDP1 led to a 40% reduction in ubiquitination activity **(Figure 2E & F)**. We conclude that SCF^bxo7^ is a major, but not the exclusive, E3 ligase recruited by Ubiquilins and that Ubiquilins can preferentially recruit the activated form of this enzyme.

### UBQLN2 shuttles substrates into biomolecular condensates

Previous work showed that polyubiquitin chains can promote UBQLN2 phase separation and that proteasomal activity is inhibited in UBQLN2 condensates, particularly with substrates marked by K63 poly-Ubiquitin chains^43,45,47,48^. We therefore hypothesized that E3 ligase recruitment by UBQLN2 protects substrates from proteasomal degradation via the formation of biomolecular condensates **(Figure 3A)**.

**Figure 3:**
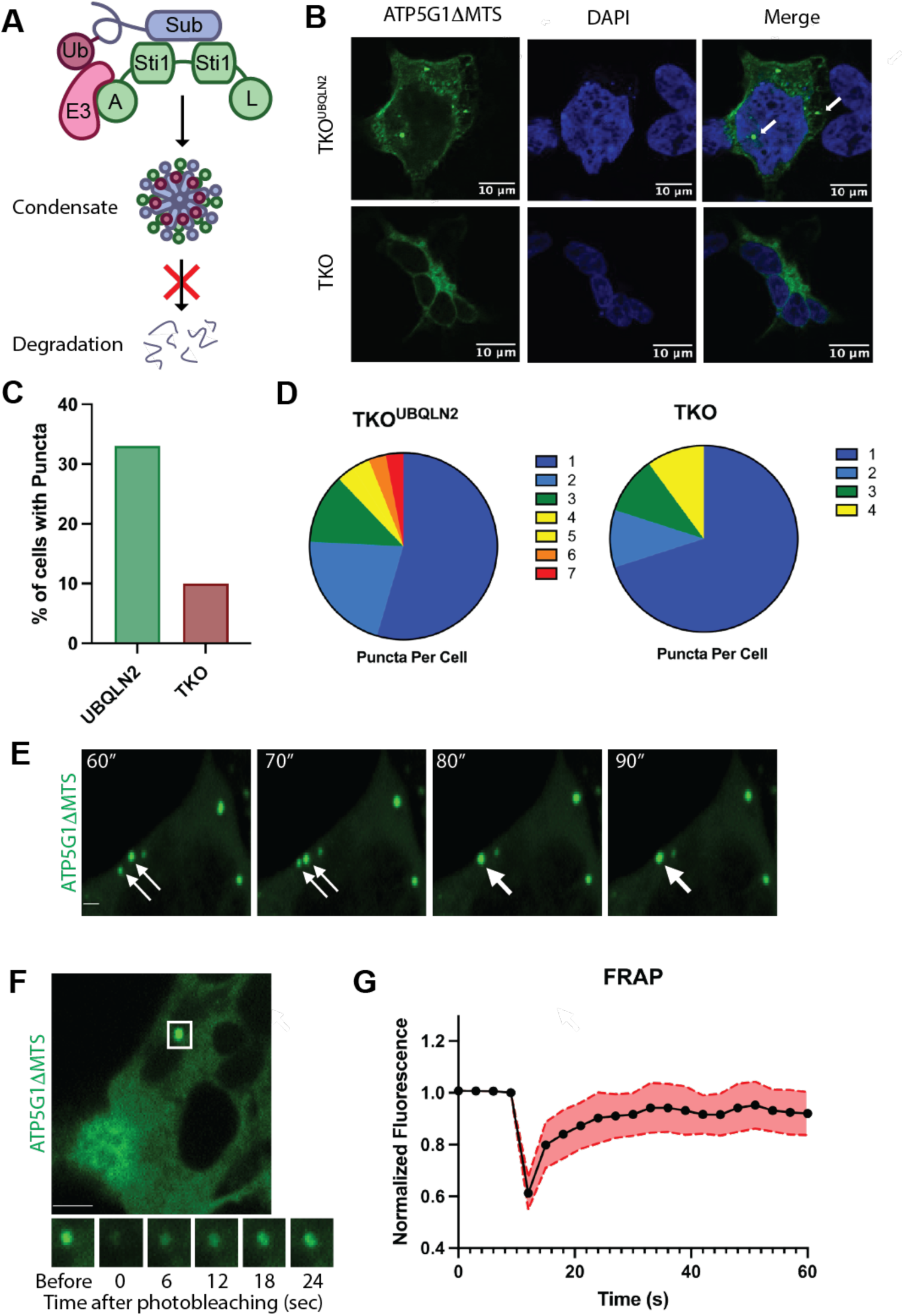
UBQLN2 shuttles substrates into biomolecular condensates. A) Proposed model for how UBQLN2 protects substrates from proteasomal degradation. UBQLN2 recruits E3 ligases via the UBA domain. Subsequent ubiquitination activity promotes formation of biomolecular condensates which protects substrates from proteasomal degradation. B) UBQLN2 promotes puncta formation of the GFP-ATP5G1ΔMTS-P2A-mCherry substrate. TKO^UBQLN2^ and TKO cells were transfected with substrate. Four hours post transfection, media was changed to include 5 ng/mL doxycycline. Cells were fixed and immunostained 24 hours post transfection. C) TKO^UBQLN2^ cells promote formation of substrate puncta. Assay in A was quantified by counting the number of substrate puncta per cell (n=100) D) TKO^UBQLN2^ cells have a higher number of puncta per cell. Pie chart showing number of puncta per cell from A. Only cells with puncta are included in the pie chart. E) Substrate puncta show liquid-like properties by droplet fusion. Substrate puncta formation was observed under oxidative stress conditions by live cell 3D imaging. See **Video S1** for full video. F) Substrate puncta show liquid-like properties with Fluorescence recovery after photobleaching (FRAP). Analysis was performed with time-lapse imaging at 3-second intervals. ROIs were chosen to analyze individual puncta in the areas with the minimal background of diffuse cytosolic fluorescence. G) Average FRAP from 10 puncta with standard error of the mean shown in red. Fluorescence intensity prior to bleaching was normalized to 1. Mobile fraction=0.534±0.135 and Half-life= 4.96±1.77 sec.

To test this hypothesis, we transfected TKO^UBQLN2^ and TKO cells with the GFP-ATP5G1ΔMTS-P2A-mCherry construct and looked for the formation of round substrate puncta. We observed a higher number of substrate puncta per cell in TKO^UBQLN2^ cells compared to the TKO cells **(Figure 3B-D)**. To test if the puncta also contained UBQLN2, we repeated the assay with anti-myc immunostaining to visualize UBQLN2. We observed strong co-localization of UBQLN2 with substrate puncta **(Fig S4A & B)**. To test if the substrate puncta also contained ubiquitin, we next co-transfected the same cell lines with the GFP-ATP5G1ΔMTS-P2A-mCherry construct and HA-Ubiquitin. Anti-HA immunostaining again shows robust co-localization of substrate puncta with Ubiquitin **(Figure S4C & D)**. Finally, to test if the puncta contain K63 specific Ubiquitin linkages, we co-transfected cells with the GFP-ATP5G1ΔMTS-P2A-mCherry construct and an HA-Ubiquitin with all lysine residues except K63 mutated to arginine. We again observed co-localization of K63 specific Ubiquitin with substrate puncta **(Figure S4E & F)**.

This prompted use to investigate if UBQLN2 phase separation promotes the formation of substrate puncta. We therefore treated TKO^UBQLN2^ cells with sodium arsenite, which was previously shown to induce UBQLN2 phase separation^23^. Upon induction of UBQLN2 expression, we observed a three-fold increase in substrate puncta formation compared to mock treated cells **(Figure S5)**.

To test if the substrate puncta have liquid-like properties, we performed 3D time-lapse live cell imaging in TKO^UBQLN2^ cells treated with sodium arsenite. We observed highly dynamic substrate puncta that undergo fusion **(Figure 3E & Movie S1)**. Finally, to monitor puncta liquidity we performed a fluorescent recovery after photobleaching (FRAP) assay and observed rapid recovery **(Figure 3F & G)**. As our puncta meet several established criteria for condensate formation^50^, we conclude that UBQLN2 promotes substrate phase separation into biomolecular condensates.

### UBQLN2 ALS mutants have defects in substrate stabilization that is exacerbated upon overexpression

The specific mechanistic defects of ALS mutations on UBQLN2 function are unknown, however many disease variants show altered phase separation properties. To better understand how ALS mutants affect UBQLN2 activity, we developed TKO rescue cell lines for the following UBQLN2 mutations, which are causative for ALS: T487I, A488T, P497L, P506T, and P506A^13^. We performed doxycycline titrations to approximate physiological expression levels **(Figure S6)** and then measured substrate degradation using the GFP-ATP5G1ΔMTS-P2A-mCherry construct. All Ubiquilin variants were able to stabilize the substrate compared to WT and TKO cells. However, none of the ALS mutants were able to stabilize substrates to the same degree as TKO^UBQLN2^ **(Figure 4A)**.

**Figure 4:**
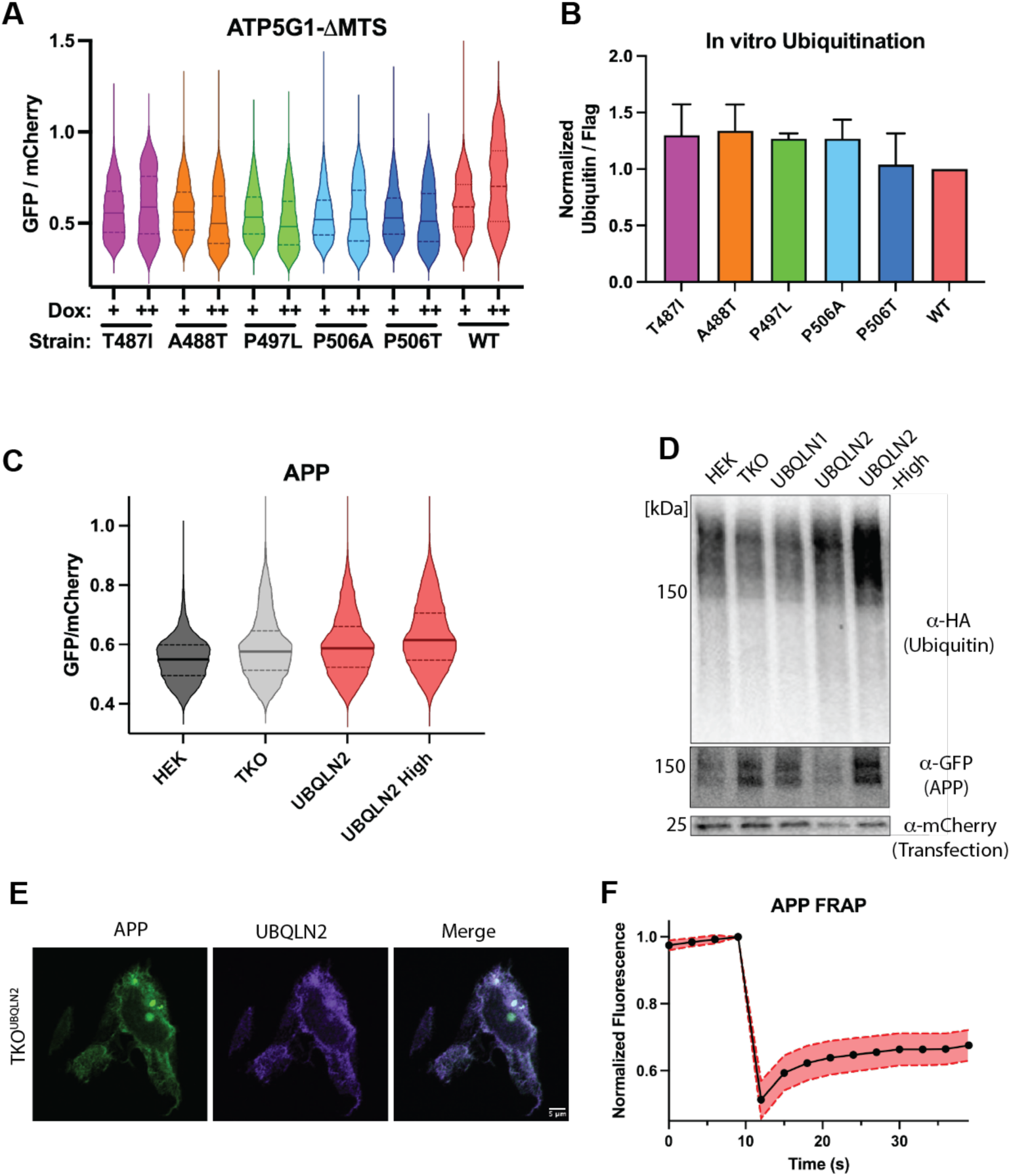
UBQLN2 ALS mutants have defects in substrate stabilization. A) UBQLN2 ALS mutants show concentration dependent defects in stabilization of model substrate ATP5G1 ΔMTS. Doxycycline was added to induce UBQLN2 expression at physiological (+) or overexpression (++) levels. B) In vitro ubiquitination assay shows no major differences in E3 ligase recruitment by UBQLN2 ALS mutants. Quantification of S6D (n = 3). Y axis is intensity of anti-ubiquitin blot divided by intensity of anti-flag blot. Data is normalized such that wild type UBQLN2 has a value of 1. C) TKO^UBQLN2^ cells stabilize APP with the degree of stabilization correlating with UBQLN2 expression levels. The GFP-APP-P2A-mCherry substrate was transfected into the indicated TKO rescue cell lines and the GFP/mCherry ratio was measured by flow cytometry. D) TKO^UBQLN2^ cells have an accumulation of ubiquitinated substrates that correlates with UBQLN2 expression levels. The GFP-APP-P2A-mCherry substrate was co-transfected with HA-Ubiquitin. GFP immunoprecipitation shows an accumulation of Ubiquitinated substrates in TKO^UBQLN2^ cells. E) APP puncta co-localize with UBQLN2. F) Average FRAP from 9 puncta with standard error of the mean shown in red. Fluorescence intensity prior to bleaching was normalized to 1. Mobile fraction=0.142±0.045 and Half-life= 4.03±1.86 sec.

Because UBQLN2 phase separation is concentration dependent, we examined the effect overexpression of the UBQLN2 ALS mutants has on substrate stabilization. Surprisingly, we observed that overexpression of most mutants led to a decrease in substrate stabilization rather than the further stabilization of substrate observed with WT UBLQN2 **(Figure 4A)**.

This prompted us to ask if the ALS mutants show a difference in their ability to recruit E3 ligases. We performed our *in vitro* ubiquitination assay with each of these ALS mutants. Total ubiquitination activity was comparable across all UBQLN2 disease mutants **(Figures 4B & S6D)**. We conclude that UBQLN2 ALS mutants have defects in substrate stabilization that are exacerbated upon UBQLN2 overexpression. The defects in substrate stabilization are not due to differences in E3 ligase recruitment, but may instead be due to different phase separation propensities of the ALS mutations.

### UBQLN2 stabilizes APP via formation of biomolecular condensates

There is a rich literature showing association of Ubiquilins with pathological aggregates that characterize proteinopathies, including Alzheimer’s disease^36,51^. Since amyloid precursor protein (APP) is a hallmark of Alzheimer’s disease and was previously shown to associate with Ubiquilins^35,36,52^, we asked if UBQLN2 can also stabilize APP via the formation of biomolecular condensates.

We first investigated if UBQLN2 can protect APP from proteasomal degradation. We cloned APP into the GFP-P2A-mCherry reporter and measured substrate stability by flow cytometry. Similar to results with ATP5G1ΔMTS, we observed substrate stabilization above the levels in TKO cells, with further stabilization upon UBQLN2 overexpression **(Figure 4C)**. To see if APP is ubiquitinated, we repeated our *in vivo* ubiquitination assay and observed an accumulation of ubiquitinated APP that increases upon UBQLN2 overexpression **(Figure 4D)**.

To test if UBQLN2 promotes the formation of APP in condensates, we transfected TKO^UBQLN2^ with the GFP-ATP5G1ΔMTS-P2A-mCherry construct. We observed co-localization of APP with UBQLN2 **(Figure 4E)**. Photobleaching of the APP puncta shows rapid recovery, suggesting that the puncta have liquid-like properties **(Figure 4F)**. Consistent with the more modest stabilization of APP, the overall mobile fraction and percent recovery were lower than with ATP5G1ΔMTS. We conclude that substrate stabilization via the formation of biomolecular condensates is a general feature of UBQLN2.

## Discussion

A hallmark of many neurodegenerative diseases is the accumulation of protein aggregates, which often arise from biomolecular condensates that have aberrant phase separation^53,54^. Interestingly, Ubiquilins frequently co-localize with these aggregates^20,32,55^. Despite decades of research, many fundamental aspects of Ubiquilin function remain unclear, including the unique role of each Ubiquilin paralog, how a proteasomal shuttle factor can paradoxically stabilize substrates, and the physiological role of Ubiquilin phase separation. Here, we have addressed these foundational questions by demonstrating that UBQLN2 has a unique ability to protect substrates from proteasomal degradation via the formation of biomolecular condensates. Importantly, when we repeated our substrate stabilization assays with UBQLN2 ALS mutants, which have altered phase separation properties, we observed defects in substrate stabilization. Overexpression of these UBQLN2 mutants exacerbated the defects in substrate stabilization, consistent with essential role of phase separation in substrate stabilization^23,26^.

We also demonstrated that UBQLN2 can recruit E3 ligases more robustly than other Ubiquilin paralogs. Because polyubiquitin chains promote UBQLN2 phase separation^44,45^, we propose that robust E3 ligase recruitment by UBQLN2 is responsible for the enhanced phase separation and subsequent substrate stabilization. This appears to be a general mechanism as we showed that UBQLN2 directly binds to APP and protects it from proteasomal degradation via the formation of biomolecular condensates.

We identified SCF^bxo7^ as one of the major E3 ligases recruited by Ubiquilins. Interestingly, Ubiquilins appear to selectively recruit the activated form of SCF^bxo7^, as removal of Nedd8, which is required for enzyme activation, leads to a ∼40% drop in ubiquitination activity. It will be important to determine the unique role of each E3 ligase and any potential deubiquitinating enzymes in regulating Ubiquilin activity. Particular focus should be on the regulated recruitment of E3 ligases that build K48- and K63-linked polyubiquitin chains, as previous work from the Castañeda lab showed that substrates with K63 polyubiquitin linkages, either alone or as mixed chains, promote condensate formation and inhibit proteasomal degradation of substrates^43^. Consistent with this model, we observed K63 ubiquitin in our substrate puncta and UBQLN2 binding to Herc1, which can build K63 polyubiquitin chains.

In summary, this study elucidates the distinct role of UBQLN2 among Ubiquilin paralogs in protecting substrates from proteasomal degradation. Counterintuitively, substrate ubiquitination reduces proteasomal degradation by promoting the formation of biomolecular condensates. This appears to be a general mechanism as the trend holds for the Alzheimer’s inducing protein APP. Together, this work provides an exciting new paradigm for understanding UBQLN2 function in health and neurodegenerative disease.

## Materials and Methods

### Cloning

The genes encoding Ubiquilins and associated variants were cloned into a pET28a vector with an N-terminal 6xHis tag followed by a 3C cleavage site and 3xFlag tag. Cloning was completed via Gibson Assembly for all Ubiquilin paralogs, disease mutants, and UBL/UBA truncations. Successful cloning was verified by Sanger sequencing.

For generation of stable cell lines, Ubiquilin variants were cloned into pcDNA5 mammalian expression vector containing an N-terminal 3xMyc tag with flanking FLP-FRT sequences for genomic integration. Cloning was completed via Gibson Assembly and restriction cloning. Successful cloning was verified by Sanger sequencing.

GFP-P2A-mCherry Dual Reporter plasmids were obtained courtesy of Manu Hedge and Sichen Shao. The genes encoding GFP-P2A-mCherry and GFP-ATP5G1ΔMTS-P2A-mCherry were cloned via restriction cloning into a pTK derivative (Thermo Fisher PI16151), under the control of the HSV TK/CMV promoter. Successful cloning was verified by Sanger sequencing.

### Protein Purification

Plasmids encoding Ubiquilin variants were transformed into *E. coli* BL21(DE3) pRIL cells and expressed in Terrific Broth at 37° C to an OD_600_ of 0.6-0.8. Cultures were induced with 0.25 mM IPTG for 4.5 hours at 16° C. Cells were harvested by centrifugation at 3900 rpm, resuspended in Ubiquilin Lysis Buffer (50 mM Hepes pH 7.5, 150 mM NaCl, 0.01 mM EDTA, 10% glycerol, 20 mM imidazole), and supplemented with 1 mM PMSF, 0.05 mg/mL lysozyme (Alfa Aesar), and 500 U of Universal Nuclease (Pierce). Cells were lysed by sonication and supernatant was isolated by centrifugation for 30 min at 12,000 rpm. Supernatant was purified by Ni-NTA affinity chromatography (Thermo Fisher) by gravity flow. Ni-NTA resin was washed with 15 column volumes of Ubiquilin Lysis Buffer and eluted with 2 column volumes of Ubiquilin Lysis Buffer supplemented with 250 mM imidazole. Protein was dialyzed overnight at 4° C against Ubiquilin Lysis Buffer supplemented with 3C Protease at a 1:100 ratio of protease to protein. Protein was passed over Ni-NTA resin to remove any uncleaved protein or Nickel binding contaminants. The flow through was collected and further purified by size exclusion chromatography on the Superdex 200 Increase 10/300 GL (GE Healthcare). Peak proteins were pooled and verified by denaturing SDS PAGE, concentrated to 60-100 μM using a 50kD MWCO centrifugal concentrator, aliquoted, flash frozen in LN2, and stored at -80° C. Protein concentration was measured by A280 and confirmed by Bradford assay.

### Cell culture

#### Maintenance

HEK23TRex cells (TKO and WT) were maintained in DMEM supplemented with 10% FBS, 100 units/mL penicillin, 100 μg/mL streptomycin, 15 μg/mL Blasticidin, and 100 μg/mL Zeocin. HEK23TRex Rescue cells were maintained in DMEM supplemented with 10% FBS, 100 units/mL penicillin, 100 μg/mL streptomycin, 15 μg/mL Blasticidin, and 150 μg/mL Hygromycin. Cells were passaged every 2-3 days once reaching a confluency of 75-90%. Cells were not used further than passage 25.

#### Cell Lysis and Normalization

Cells were washed with DPBS, Trypsinized and centrifuged at 100g for 5 min. Cell pellets were lysed by mixing with the lysis buffer (25 mM Tris-HCl, pH 7.4, 150 mM NaCl, 1mM EDTA, 1% NP-40 and 5% Glycerol) supplemented with 125 Units/mL of universal nuclease (Pierce) and 1x Halt™ Protease Inhibitor Cocktail (Thermo Scientific). Lysates were centrifuged at 14,000 rpm for 15 min at 4° C and the supernatant was normalized by A280 with cell lysate dilution buffer (25 mM Tris-HCl, pH 7.4, 150 mM NaCl, 0.5 mM EDTA).

#### Induction of Ubiquilin expression

One day (24 h) prior to doxycycline addition, cells were split into 10 cm dishes in DMEM media and 10% FBS lacking antibiotics. Ubiquilin expression was induced by addition of doxycycline according to **Table 1**. After 24 h in media with doxycycline, cells were harvested.

**Table 1:** Doxycycline concentrations used to induce Ubiquilin expression.

| Cell line | Physiological expression | Overexpression |
| --- | --- | --- |
| TKO<sup>UBQLN1</sup> | 5 ng/mL | n/a |
| TKO<sup>UBQLN2</sup> | 0.4 ng/mL | 5 ng/mL |
| TKO<sup>UBQLN4</sup> | 5 ng/mL | n/a |
| TKO<sup>T487I</sup> | 0.25 ng/mL | 5 ng/mL |
| TKO<sup>A488T</sup> | 1 ng/mL | 5 ng/mL |
| TKO<sup>P497L</sup> | 3 ng/mL | 5 ng/mL |
| TKO<sup>P506A</sup> | 2.5 ng/mL | 5 ng/mL |
| TKO<sup>P506T</sup> | 2 ng/mL | 5 ng/mL |

#### Transfection

Transient transfection was conducted in 10 cm culture dishes. 24 hours prior to transfection, HEK293TRex cells were split into 10 cm dishes with DMEM supplemented with 10% FBS but lacking antibiotics. 36 μg of DNA was transfected into cells using 18 μL Lipofectamine 2000 (Invitrogen) in Opti-MEM (Gibco). For co-transfections with HA-Ubiquitin, 7.5 μg of the substrate and 4.5 μg of HA-Ubiquitin were transfected with 30 μL Lipofectamine 2000 (Invitrogen) in Opti-MEM (Gibco). Opti-MEM media was exchanged for DMEM supplemented with 10% FBS but lacking antibiotics 4-5 hours following transfection. For experiments in which Ubiquilins were induced with Doxycycline, the media for the media change was supplemented with doxycycline according to **Table 1**. Cells were harvested for subsequent analysis or experiment 24 hours following transfection.

#### Generation and validation of TKO rescue cell lines

TKO HEK293TRex cells were obtained courtesy of Manu Hegde and were generated as described previously^32^. TKO cells were transfected, as described above, in 6 well plates. 0.6 μg 3xMyc-Ubiquilin was co-transfected with 5.4 μg pOG44 at a 9:1 ratio of pOG44 to plasmid in Opti-MEM (Gibco) using 3 μL Lipofectamine 2000 (Invitrogen). Opti-MEM media was exchanged for DMEM supplemented with 10% FBS but lacking antibiotics for 4-5 hours following transfection. 24 hours following transfection, media was exchanged for fresh DMEM with 10% FBS. 48 hours following transfection, cells were split into 10 cm culture dishes and cultured in DMEM supplemented with 10% FBS, 100 units/mL penicillin, 100 μg/mL streptomycin, 15 μg/mL Blasticidin, and 150 μg/mL Hygromycin (Gibco). Cells were frozen following 17-21 days of antibiotic selection. Expression of transfected species was verified by western blots (anti-Myc and anti-Ubiquilin) as well as PCR amplification and sequencing.

### Immunoprecipitation

#### From RRL

30 μL of anti-Flag magnetic resin (Pierce) was washed in Flag Pulldown Buffer (25 mM HEPES, 250 mM KOAc, 0.1% Tween 20). RRL containing Flag-tagged Ubiquilin was incubated with resin at 4° C for 1 hour. Flow through was discarded, and resin was further washed in Flag Pulldown Buffer. Bound proteins were eluted with 1 mg/mL Flag Peptide (ApexBio) at 37° C on thermomixer.

#### From cells

Magnetic resin (anti-GFP, Proteintech) was washed in Cell Lysate IP Wash Buffer (25 mM Tris-HCl, 150 mM NaCl, 0.5 mM EDTA, 0.1% Tween 20). Cell Lysates were incubated with resin at 4° C for 1 hour. Flow through was discarded, and resin was further washed with Cell Lysate IP Wash Buffer. For anti-GFP resin, bound proteins were eluted with 1x SDS buffer at 95° C for 5 minutes on a thermomixer.

### Cell Viability Assay

Cells were maintained in 6 well plates with the growth conditions as described and counted every 24 hours from seeding for 4 consecutive days. Cells were harvested and pelleted as described and the pellets were dissolved in DPBS. The cell suspension was mixed with 4% Trypan blue (Gibco) at a 1:1 ratio and the cells were counted using a hemocytometer.

### Flow Cytometry Degradation Assays

HEK293TRex Cell Lines were transfected as described above in 10 cm dishes with GFP-P2A-mCherry dual reporter plasmids. TKO rescue cell lines were induced with indicated levels of doxycycline 4-5 hours following transfection. 24 hours following transfection, cells were washed in 10 mL DPBS, trypsinized, and centrifuged at 100 x g for 5 minutes. Cells were resuspended in DPBS and filtered via 40/50 um filter into 5 mL tubes. Cells were stained in 3 μM DAPI (Invitrogen). Flow Cytometry was completed on BioRad ZE5 Cell Analyzer or BD Biosciences LSR15 cytometer, and further analysis was completed using FlowJo.

### Ubiquitination Assay

#### RRL IVT

RRL *in vitro* transcription and translation was performed as described previously^56^. DEPC water was used throughout the protocol. Omp25 IVT optimized plasmid was transcribed via T7 polymerase in the presence of RNase inhibitor (Invitrogen) for 1 hour at 37° C. Translation was carried out at 32° C for 30 minutes in the presence of recombinantly expressed, purified Ubiquilins at a concentration of 10.4 μM. Translation was supplemented with 5 μM methionine.

#### In vitro Ubiquitination

Ubiquitination of IVT reaction products was performed as described previously^32^. 4x Ubiquitination Mix was assembled using 40 μM His-ubiquitin (R&D Systems), 0.4 μM GST-Ube1 (R&D Systems), 1.2 μM UbcH5a (R&D Systems), and an energy regenerating system (4 mM ATP (Thermo Scientific), 40 mM Creatine Phosphate (Thermo Scientific), and 160 μg/mL Creatine Kinase. Ubiquitination was performed for 1 hour at 37° C on thermomixer, followed by termination in 1% SDS at 95C for 10 minutes on thermomixer.

#### NEDP1

For experiments in which NEDP1 treatment was used, RRL was incubated with 10 μM His-NEDP1 (Southbay Bio) at 37° C for 30 minutes on thermomixer. Treated RRL was used in subsequent ubiquitination or immunoprecipitation experiments.

### Western blots

SDS PAGE was performed using 4% stacking, 15% separating Tris-Glycine gels (cast in house) (Figure 2B), 4–20% Mini-PROTEAN TGX Precast Protein Gels (Bio-Rad), 4–20% Criterion TGX Stain-Free Protein Gels (Bio-Rad). Gels were transferred to 0.45 um PVDF transfer membrane (Thermo Scientific) at 4° C for 1 hour at 330 mA in 1x Transfer Buffer (192 mM Glycine, 25 mM Tris-Base, 0.01% SDS, in MeOH) or using the Trans-Blot Turbo Transfer System (BioRad) at mixed molecular weight settings for 10 minutes. Blocking was completed in 5% milk, antibody incubations in 1% milk, and washing in 1x TBST (20 mM Tris-Base, 200 mM NaCl, 0.1% Tween-20, pH 7.4), all using standard techniques. Antibody dilutions can be found in the Key Resources Table below. Blots were developed using Cytiva Amersham ECL Select Western Blotting Detection Reagent (Cytiva) or SuperSignal West Femto Maximum Sensitivity Substrate (Thermo Scientific).

### Immunofluorescence

Cells grown and transfected on uncoated glass coverslips (Electron Microscopy Sciences), fixed with 2% paraformaldehyde for 20 min and washed 2 times with PBS. Cells were permeabilized with 0.1% Triton X-100 for 20 min and blocked with 5% goat serum with PBB (0.5% BSA in PBS). Cells were washed 5 times with PBB and incubated with the primary antibody at RT for 1 hour. Cells were washed 6-7 times with PBB and incubated with the secondary antibody at RT for 1 hour. Cells were washed 5 times with PBB following 5 washes with PBS. Next cells were incubated with 1mg/100mL Hoechst stain for 1 min followed by 3 PBS washes with 1 min incubations in between. Coverslips were adhered onto a glass slide using Gelvatol. Slides were cured for 24 hours and imaged.

Slides were imaged with a Nikon A1 confocal system with a 60x objective and 1.4 NA. Image analysis was done using a macro created by Fiji for thresholding and particle analysis using an intensity cutoff of 2500, a size cutoff of 0.07 micron^2^, and a circularity cutoff of 0.75 for the unstressed conditions. Stressed condition puncta were analyzed with an intensity cutoff of 700, a size cutoff of 0.06 micron^2^, and no circularity cutoff.

### Live-cell confocal microscopy

Cells were seeded on 35 mm MatTek dishes with a 14 mm microwell and transfected with 1 μg of DNA and 3 μL of Lipofectamine 2000 (Invitrogen) in Opti-MEM (Gibco). Opti-MEM media was exchanged for DMEM supplemented with 10% FBS, but lacking antibiotics, for 4 hours following transfection. Doxycycline induction of Ubiquilin expression was performed during the media exchange 4 hours post transfection using the concentrations listed in **Table 1**. Cells were imaged 20-24 hours post transfection.

Live-cell imaging was performed using Marianas spinning disk confocal imaging system workstation based on a Zeiss Axio Observer Z1 inverted fluorescence microscope equipped with 63x Plan Apo PH NA 1.4 objective, Spherical Aberration Correction unit, Yokogawa CSU-W1, Vector photomanipulation module, Andor iXon EMCCD camera, environmental chamber, piezo stage controller and lasers (405, 445, 488, 515, 561, and 640 nm), all controlled by SlideBook24 software (Intelligent Imaging Innovation, Denver, CO).

To perform time-lapse three-dimensional (3D) imaging, z-stacks of confocal images (20 serial two-dimensional confocal images at 400 nm z-steps) were acquired through the 488 nm channel from cells grown on 35 mm Mat-Tek dishes every 5 seconds at 37°C in 5% CO_2_.

To perform fluorescence recovery after photobleaching (FRAP) analysis, time-lapse images were collected at 3-second intervals from a single confocal plane through the 488 nm channel. Photobleaching was performed on user-selected regions of interest (ROIs; 2X2 μm squares) using Vector photomanipulation module. ROIs were chosen to analyze individual puncta in the areas with the minimal background of diffuse cytosolic fluorescence. ROIs from time-lapse images with stable focus and minimal lateral movement of the puncta were analyzed using the “FRAP” module in SlideBook24 software to obtain values of t_D_ (recovery half-time) and mobile fraction (M_f_, or fraction recovered). Image-wide photobleaching was negligible during the duration of the experiment under image acquisition conditions used, as confirmed by measuring the fluorescence intensity of an ROI in cells that were not photobleached through the Vector module.

### Mass spectrometry and data analysis

Mass spectrometry (MS) was performed at the University of Michigan chemistry mass spectrometry facility. 10.4 μM flag tagged, WT or ΔUBL UBQLN1 were added to 41 μL of hemin and nuclease treated RRL, incubated for 30 minutes at 32° C, and isolated by anti-flag IP. Elution from anti-Flag IP were prepared for mass spectrometry via disulfide reduction, aklylation, and trypsin digestion at 38° C overnight. Digested peptides were separated on a C18 column (Acclaim PepMap 100 C18 HPLC column, #164540) and analyzed on the nanoUPLC (Ultimate 3500) linked to Orbitrap Fusion Lumos Tribid MS. Protein identification was performed by Higher Energy Collisional Dissociation MS-MS in Data Dependent Acquisition mode and then analyzed with Protein Discoverer (v 2.2) against UniProt-Rabbit (Oryctolagus cuniculus) database. Common contaminants, including human keratin and trypsin were removed. M-oxidation, protein N-terminal acetylation, and fixed C-cabamidomethylation were included in the search. A false discovery rate filter of 1% was applied for peptide and protein IDs.

### Key Resources Table

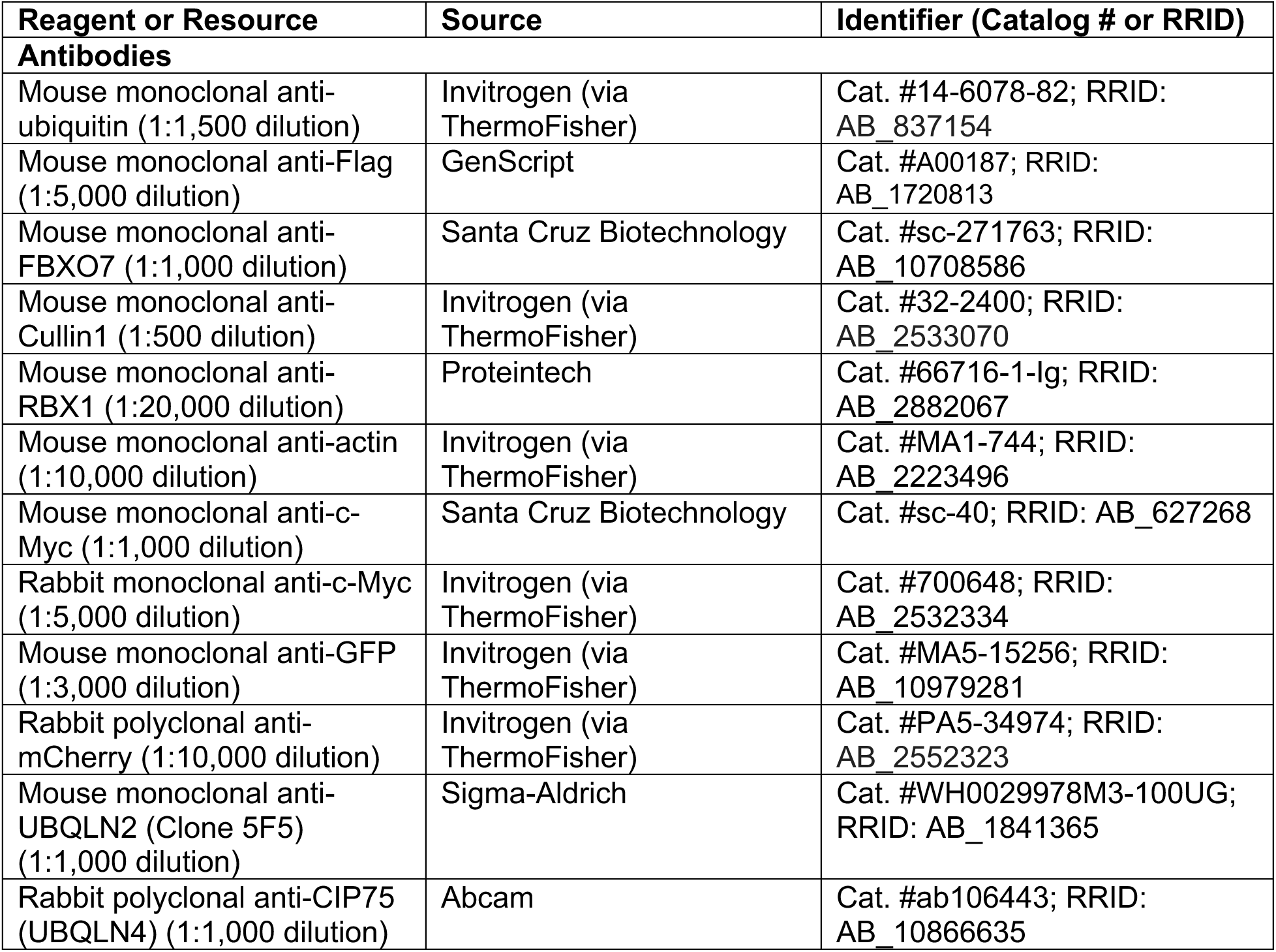
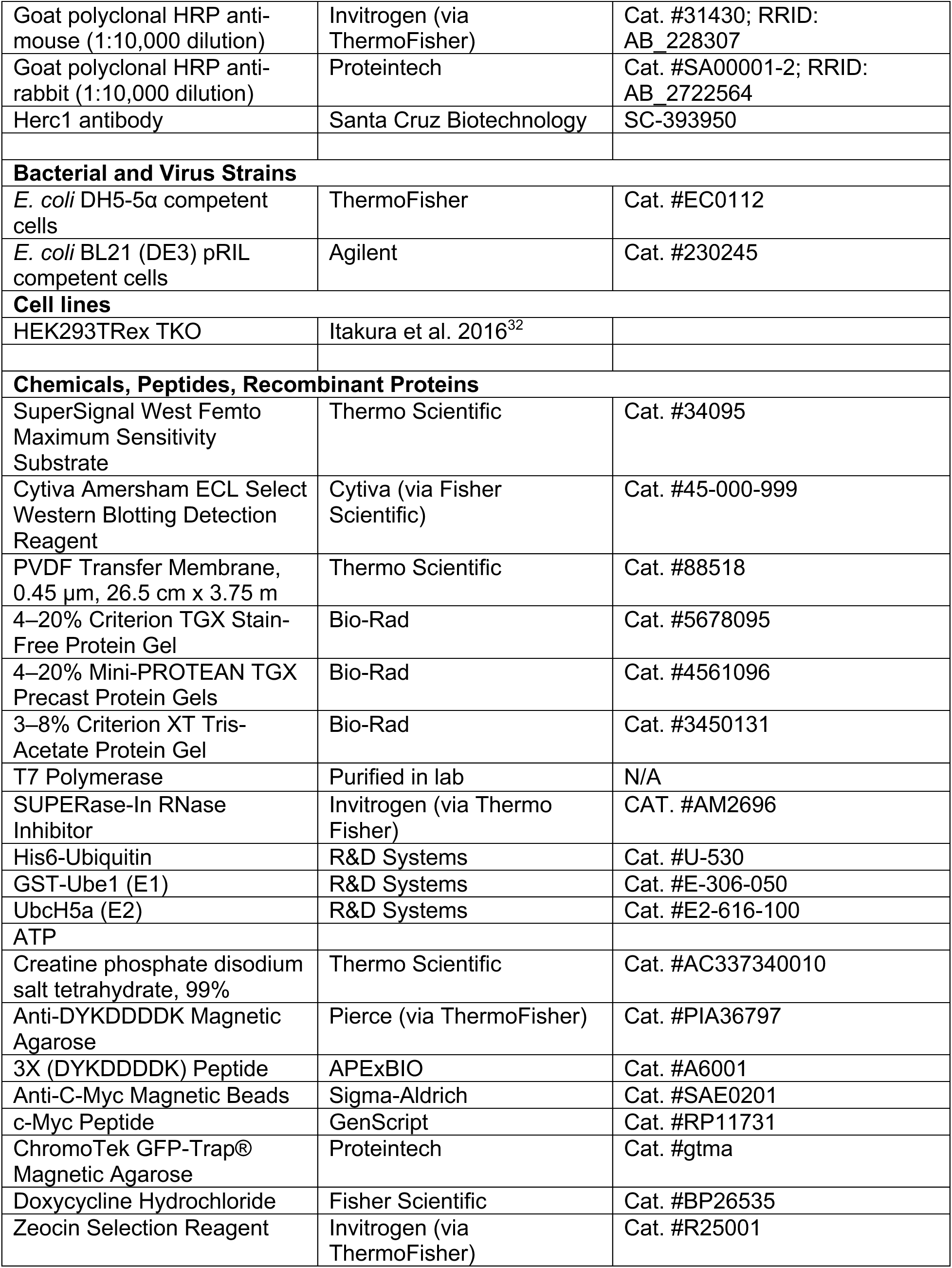
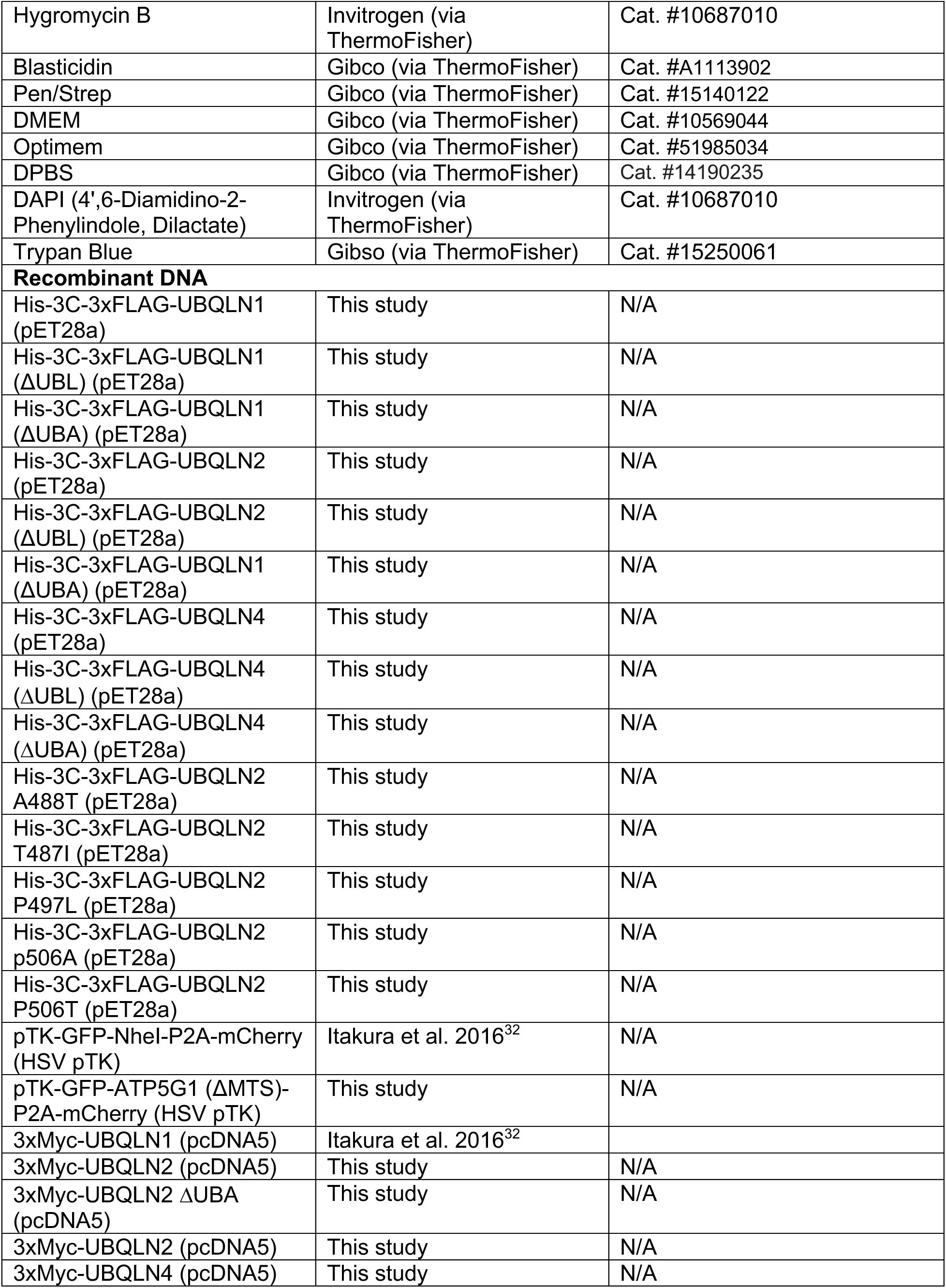
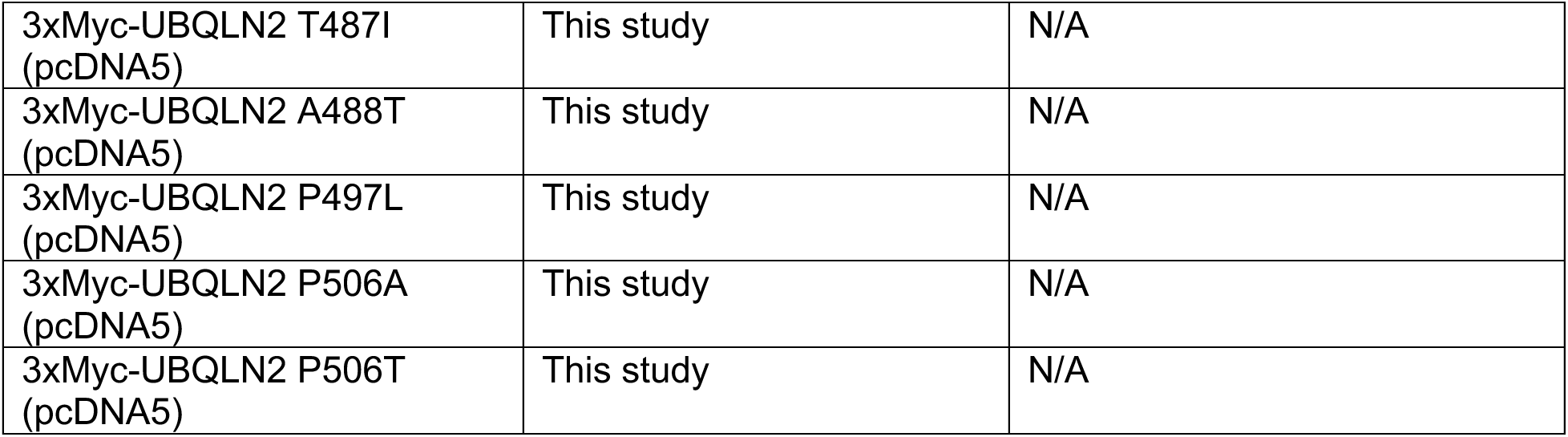

## Data availability

All data is available upon request.

## Acknowledgements

The authors wish to thank the Hegde, Shao, O’Donnell, and Castañeda labs for the TKO knockout cell line, plasmids, and helpful discussions. The authors also acknowledge Shawn Allen for help in maintaining the cell lines and the flow cytometry cores at the University of Michigan and University of Pittsburgh. Mass spectrometry data collection and analysis was supported by the Office of the Director, National Institutes of Health award number S10OD021619.

## Funding

This work was supported by NSF CAREER Award 2343131 (MLW).

## Author Contributions

Conceptualization: ANS, ST, GS, ADS, MLW; Data curation: ANS, ST, GS, ADS, MLW; Formal analysis: ANS, ST, GS, ADS, MLW; Funding acquisition: MLW; Investigation: ANS, ST, GS, ADS; Methodology: ANS, ST, GS, ADS; Project administration: MLW; Resources: ANS, ST, GS, ADS, MLW; Software: Not applicable; Supervision: MLW; Validation: ANS, ST, GS, ADS; Visualization: ANS, ST, GS, ADS, MLW; Writing – original draft: ANS, MLW; Writing – review & editing: ANS, ST, GS, ADS, MLW;

## Competing Interests

The authors declare no competing interests.

## Supplemental Figures

**Figure S1:**
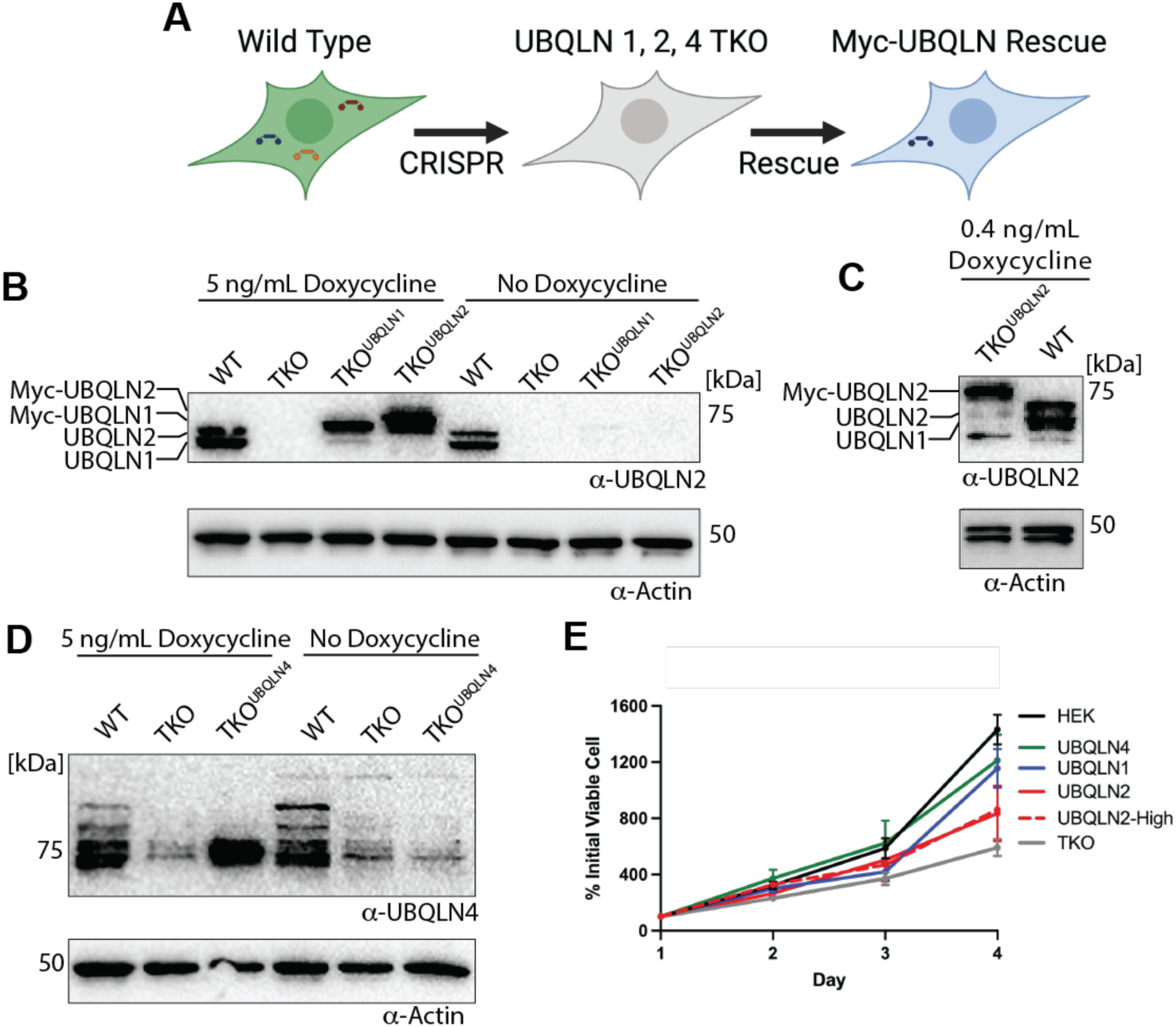
Development of TKO rescue cell lines with physiological and overexpression of individual ubiquilins. A) Diagram of TKO rescue strategy. B) TKO^UBQLN^^1^ cells require 5 ng/mL of doxycycline to have physiological Ubiquilin expression levels. TKO^UBQLN2^ cells show overexpression under these conditions. Anti-UBQLN2 western blot of cells treated with 5 ng/mL doxycycline. Anti-UBQLN2 antibody demonstrates cross-reactivity with UBQLN1^32^. C) TKO^UBQLN2^ cells require 0.4 ng/mL of doxycycline to have physiological Ubiquilin expression level. Anti-UBQLN2 western blot of cells treated with 0.4 ng/mL doxycycline D) TKO^UBQLN4^ cells require 5 ng/mL of doxycycline to have physiological Ubiquilin expression levels. Anti-UBQLN4 western blot with 5 ng/mL doxycycline E) Growth curves of WT, TKO, TKO^UBQLN1^, TKO^UBQLN2^, TKO^UBQLN4^, TKO^UBQLN2-High^ cells. Cell viability was measured with a hemocytometer and trypan blue staining.

**Figure S2:**
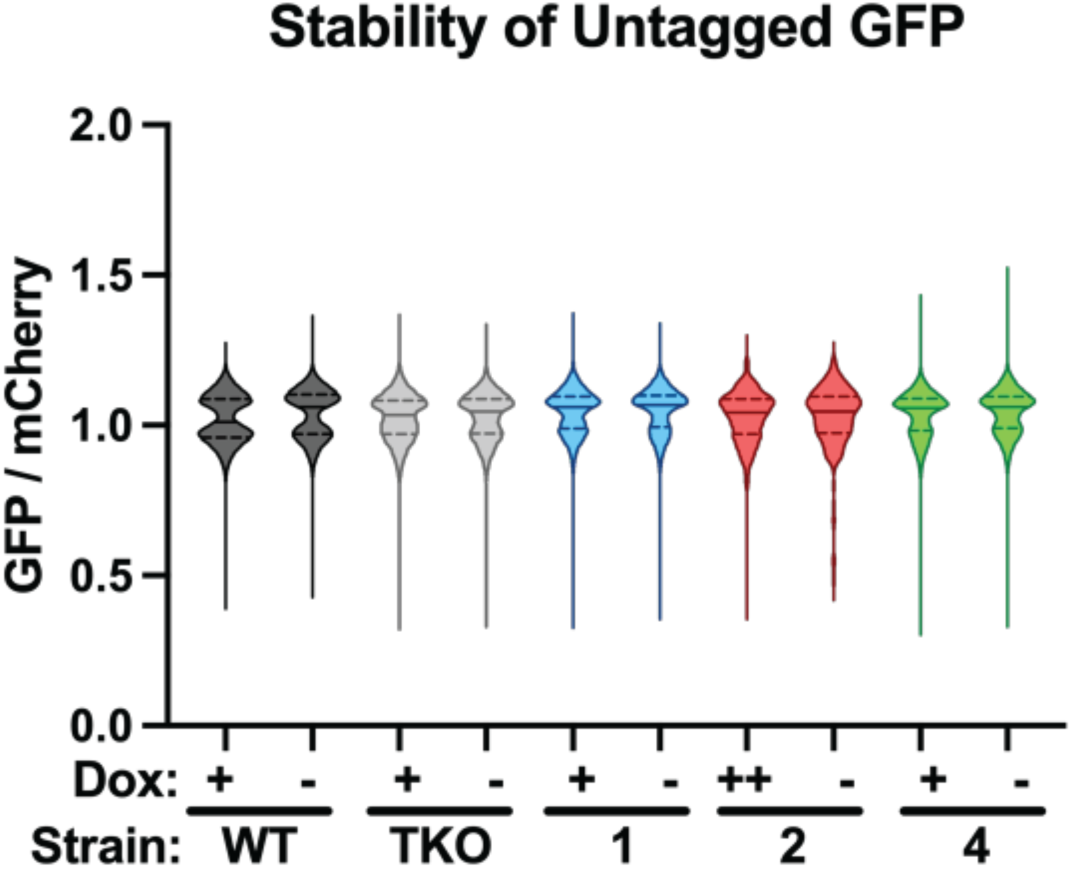
Untagged control for GFP:mCherry assy. Untagged GFP and mCherry are equally stable in all cell lines. Doxycycline concentrations were sufficient to give physiological Ubiquilin expression (+) or overexpression (++).

**Figure S3:**
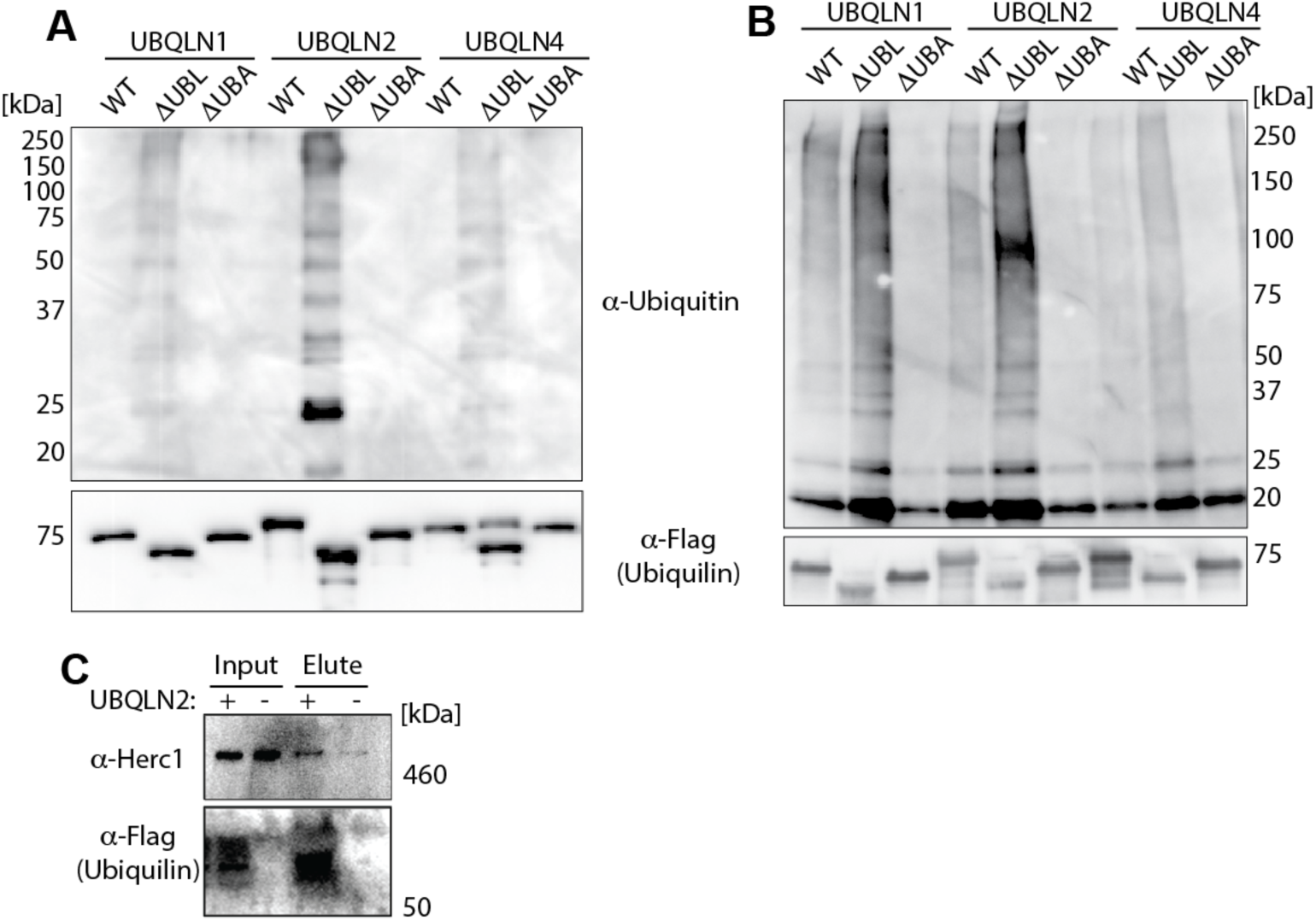
Additional replicates of the *in vitro* ubiquitination assay. A-B) *In vitro* ubiquitination assay was performed as in Figure 2. Note that replicates were performed with different stocks of purified Ubiquilins. C) ΔUBL UBQLN2 interacts with Herc1. Assay was performed as in Figure 2C, except 30 μM UBQLN2 was mixed with 100 μL of RRL.

**Figure S4:**
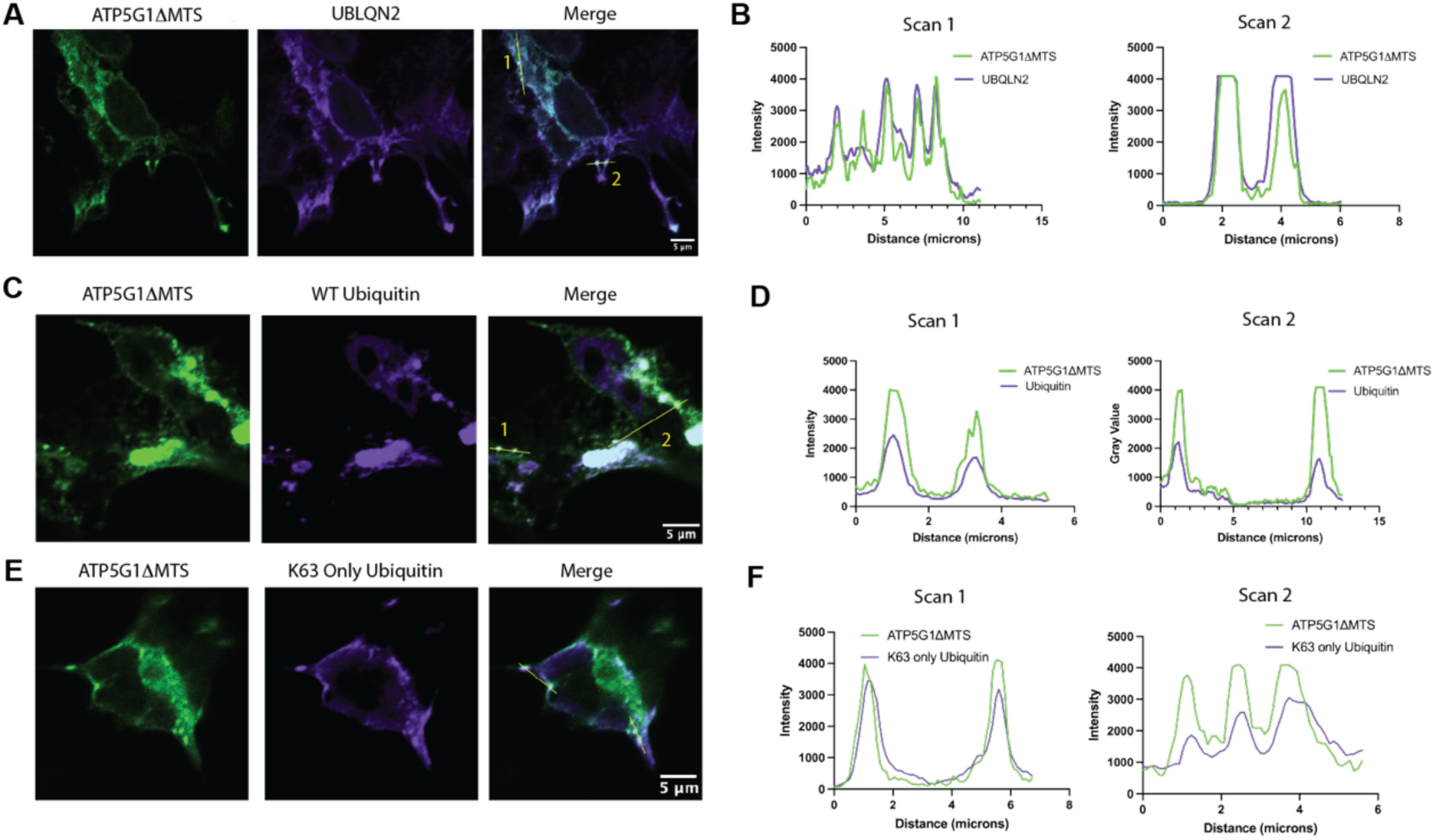
Substrate puncta co-localize with UBLQN2 and Ubiquitin. A) UBQLN2 is present in substrate puncta. TKO^UBQLN2^ cells were transfected with substrate. Four hours later, samples were treated with 5 ng/mL doxycycline for 20 h, and then fixed observed by anti-Myc immunofluorescence. B) Line scan of A shows co-localization of UBLQN2 and substrate puncta. C) Ubiquitin is present in substrate puncta. Assay performed as in A, except the substrate was co-transfected with HA-Ubiquitin and Ubiquitin was observed by anti-HA immunofluorescence. D) Line scan of C shows co-localization of Ubiquitin and substrate puncta. E) Substrate puncta contain K63 ubiquitin. Assay performed as in C, except cells were transfected with K63-only HA-Ubiquitin. F) Line scan of E shows co-localization of K63-Only Ubiquitin and substrate puncta.

**Figure S5:**
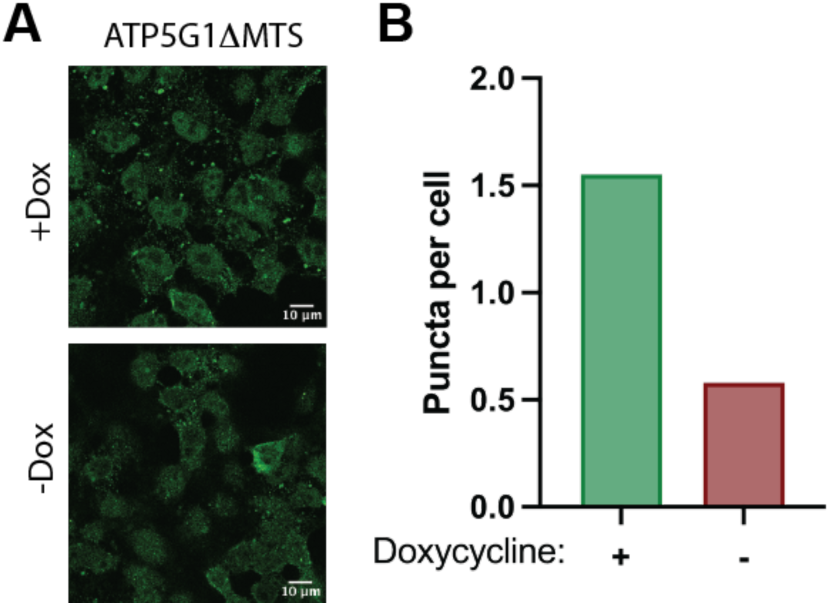
UBQLN2 promotes substrate puncta formation under oxidative stress. A) UBQLN2 expression promotes substrate puncta formation. TKO^UBQLN2^ cells were transfected with GFP-ATP5G1ΔMTS-P2A-mCherry substrate. Four hours post transfection, media was changed to include 5 ng/mL doxycycline for an additional 20 h. Cells were treated with 500 μM Sodium Arsenite for 30 minutes prior to fixation and immunofluorescence imaging. B) Quantification of data in A.

**Figure S6:**
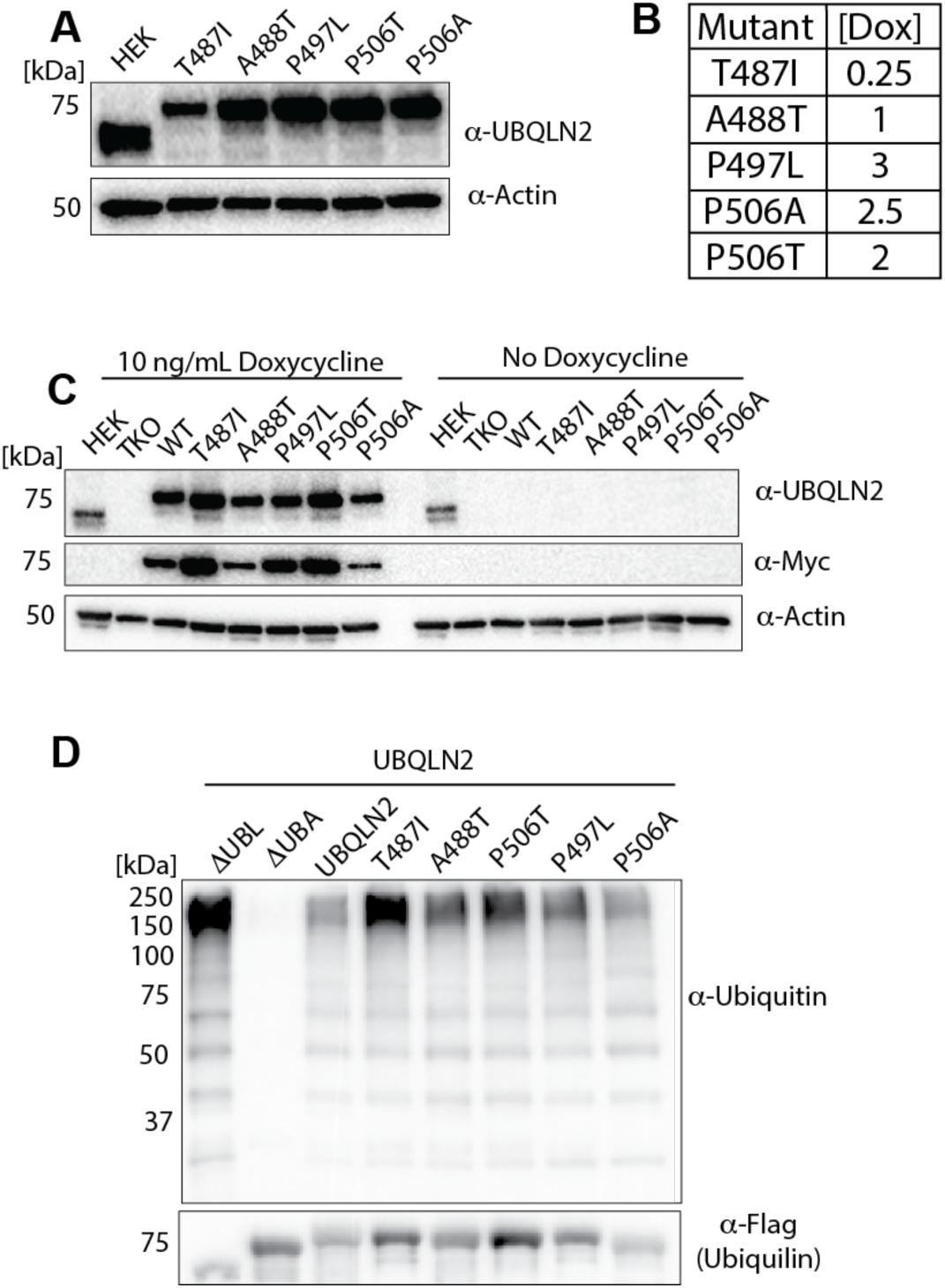
Characterization of UBQLN2 ALS Mutants. A) Anti-UBQLN2 western blot showing roughly physiological expression of ALS mutants under physiological conditions. B) Table showing the concentration of doxycycline (ng/mL) used in A. C) Anti-UBQLN2 and anti-Myc western blots of ALS mutants under overexpression conditions (10 ng/mL doxycycline) D) Representative data for in vitro ubiquitination assay with UBQLN2 ALS mutants.

**Table S1: Mass spectrometry results with full-length UBQLN1**

**Table S2: Mass spectrometry results with ΔUBL UBQLN1**

**Movie S1: Substrate puncta undergo fusion in TKO^UBQLN2^ cells** Stills for Figure 3E were captured from this movie.

## Notes

### Competing Interest Statement

The authors have declared no competing interest.

### Summary of Updates

We have performed extensive microscopy analysis to show that UBQLN2 promotes substrate phase separation. Furthermore, we show that our model holds for multiple substrates, including amyloid precursor protein.

